# Polygenic patterns of adaptive introgression in modern humans are mainly shaped by response to pathogens

**DOI:** 10.1101/732958

**Authors:** Alexandre Gouy, Laurent Excoffier

## Abstract

Anatomically modern humans carry many introgressed variants from other hominins in their genomes. Some of them affect their phenotype and can thus be negatively or positively selected. Several individual genes have been proposed to be the subject of adaptive introgression, but the possibility of polygenic adaptive introgression has not been extensively investigated yet. In this study, we analyze archaic introgression maps with refined functional enrichment methods to find signals of polygenic adaptation of introgressed variants. We first apply a method to detect sets of connected genes (sub-networks) within biological pathways that present higher-than-expected levels of archaic introgression. We then introduce and apply a new statistical test to distinguish between epistatic and independent selection in gene sets of present-day humans. We identify several known targets of adaptive introgression, and we show that they belong to larger networks of introgressed genes. After correction for genetic linkage, we find that signals of polygenic adaptation are mostly explained by independent and potentially sequential selection episodes. However, we also find some gene sets where introgressed variants present significant signals of epistatic selection. Our results confirm that archaic introgression has facilitated local adaptation, especially in immunity-related and metabolic functions and highlight its involvement in a coordinated response to pathogens out of Africa.

## Introduction

As anatomically modern humans expanded out of Africa, they encountered other groups of now extinct humans such as Neandertals and Denisovans. The availability of complete genomes from Neandertal and Denisovan individuals led to the discovery of past gene flow between these archaic hominins and non-Africans (Green, et al. 2010; Prüfer, et al. 2014; Prüfer, et al. 2017). Indeed, estimates of Neandertal ancestry for non-African populations range from 1 to 3% (Green, et al. 2010; Prüfer, et al. 2014; Prüfer, et al. 2017), and Melanesians and East Asians have an additional 3-4% and 0.2% ancestry, respectively, from potentially different sub-groups of Denisovans (Reich, et al. 2010; Meyer, et al. 2012; Prüfer, et al. 2014; Vernot, et al. 2016; Prüfer, et al. 2017; Browning, et al. 2018).

Recent statistical developments have made possible the identification of the portions of the human genome that were introgressed and survive in present-day individuals (Prüfer, et al. 2014; Sankararaman, et al. 2014; Vernot and Akey 2014; Sankararaman, et al. 2016; Vernot, et al. 2016). Many of these archaic variants encompass protein-coding genes and other functional genomic features, potentially impacting phenotypes. Indeed, the presence of archaic variants in modern humans can affect patterns of gene expression (Dannemann, et al. 2017; McCoy, et al. 2017) and influence various phenotypes, including for example immune and pigmentation traits (Gittelman, et al. 2016; Simonti, et al. 2016). Consequently, we expect some archaic variants to have an impact on the fitness of modern humans, and therefore to be the target of different forms of selection. The significant depletion of introgressed archaic variants in coding sequences has been attributed to negative selection (Sankararaman, et al. 2014; Vernot and Akey 2014; Harris and Nielsen 2016; Juric, et al. 2016), but several studies have also described cases of adaptive introgression, i.e. when a variant of archaic origin is positively selected after the introgression event. For example, the introgression of a Denisovan haplotype in EPAS1 could have helped Tibetans adapting to high-altitude (Huerta-Sanchez, et al. 2014) and single-locus studies have identified adaptive introgression in genes involved in immunity and metabolism, e.g. genes of the major histocompatibility locus (MHC) (Abi-Rached, et al. 2011), *SLC16A11* (SIGMA Type 2 Diabetes Consortium 2014), *OAS1, STAT2*, or Toll-like receptors (Deschamps, et al. 2016; Quach, et al. 2016).

Previous studies of adaptive introgression have mainly focused on isolated genes or clusters of physically linked genes yielding a strong signal of selection. However, adaptation can sometimes result from subtle shifts in allele frequencies and some traits have a polygenic basis (Stephan 2016). These faint signals would typically remain below the detection threshold of most methods and evidencing polygenic trait evolution thus remains challenging (Le Corre and Kremer 2012). Nevertheless, methods based on gene set enrichment (Daub, et al. 2013) or network analysis (Gouy, et al. 2017) have successfully detected more subtle signals of selection and identified biological pathways underlying adaptation. However, it is still difficult to distinguish between two possible modes of polygenic selection when analyzing genomic data. Observed changes in allele frequencies at different loci that can indeed be either due to i) independent selection events having sequentially occurred at different times on different genes or ii) simultaneous selection of combination of variants in different genes. We will hereafter refer to these two types of multigenic selection as *independent* and *epistatic* selection, respectively.

In this study, we analyze archaic introgression maps that have been recently made available for 35 Melanesian individuals as well as samples from the 1000 Genomes project (1000 Genomes Project Consortium 2015). We used recent and refined functional enrichment methods based on biological pathway analysis. First, we apply a method to detect sets of connected genes (sub-networks) within pathways that present higher-than-expected levels of archaic introgression. We also introduce a new test to distinguish between the sequential selection of introgressed alleles at different loci and epistatic selection by testing whether some introgressed variants are co-segregating within individuals.

## Results

### Detection of gene subnetworks enriched in archaic variants

We first computed an archaic introgression score for each gene as the density of archaic haplotypes along the exons of a given gene (see Methods). To see if groups of functionally related genes presented significantly high levels of introgression, we then used the R package *signet* (Gouy, et al. 2017) to identify high scoring subnetworks among three biological pathway databases (KEGG, NCI, and Reactome), and this for each of the three populations considered (East Asians: EAS; Europeans: EUR; Papua New Guineans: PNG). Significant pathways and associated subnetworks are presented in Table 1. As several subnetworks overlapped (and were sometimes identical), we merged all significant subnetworks for each population and assigned them to clusters (Table 1).

**Table 1:**
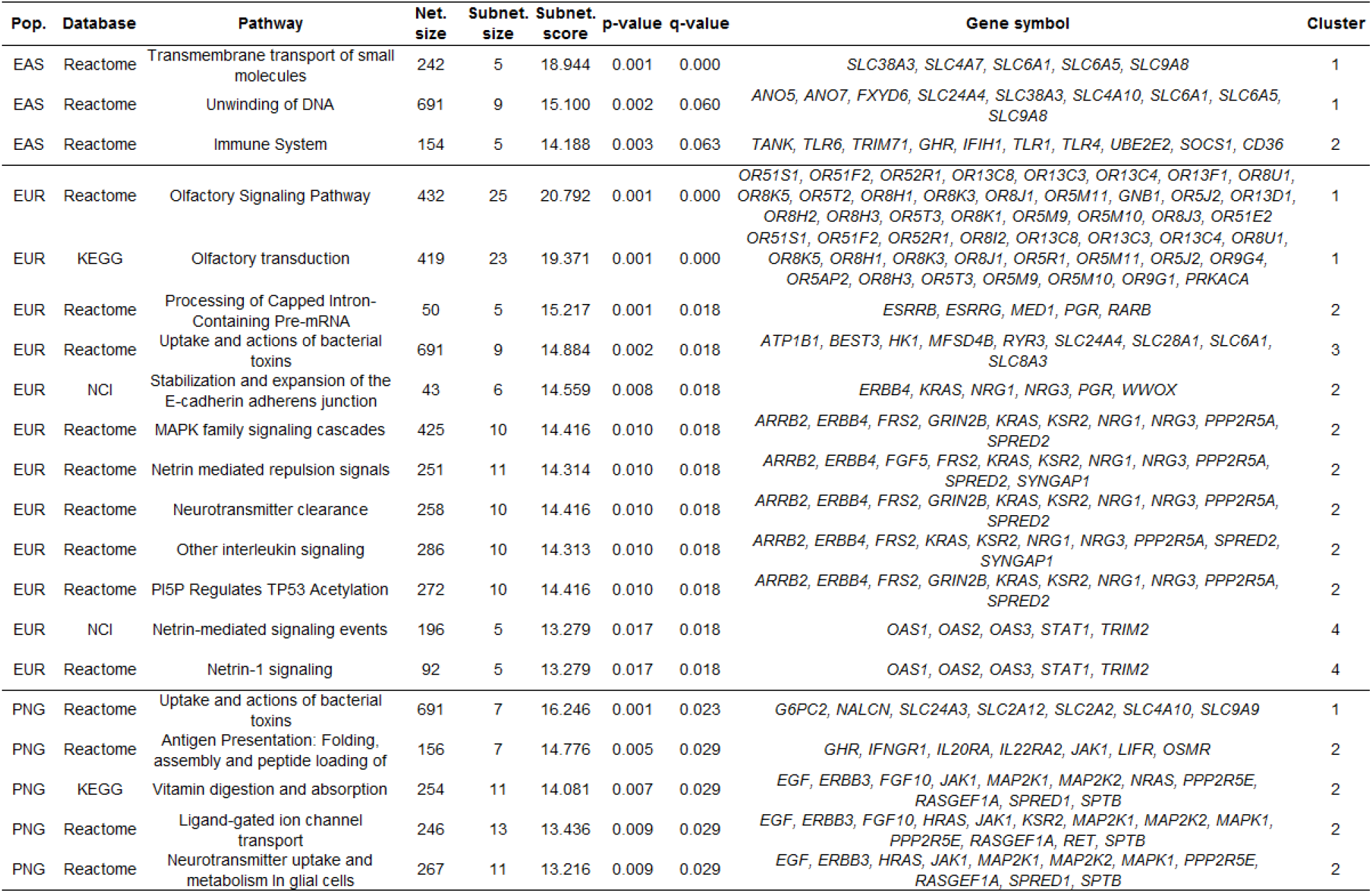
Gene subnetworks with significantly (FDR<0.05) high introgression scores in three tested population (East Asia: EAS; Europe: EUR, and Papua-New-Guinea: PNG). For each significant pathway, we report the database it belongs to, the full pathway name and sizes, the subnetwork size, its score, its p-values and its q-values, as well as the symbols of the genes belong to the significant subnetwork. Subnetworks have been assigned to the same cluster if they were overlapping for at least one gene.

In East Asians (EAS), we identify three significant pathways belonging to two non-overlapping clusters of genes. Cluster 1: “Transmembrane Transport of Small Molecules” and “Unwinding of DNA”; Cluster 2: “Immune System” (Table 1). Interestingly, several genes of the two pathways of cluster 1 belong to the Solute Carrier family (SLC). Even though *SLC* proteins are all transporters, they are involved in a variety of distinct functions. Some of them are involved in neurotransmission in the γ-aminobutyric acid (GABA) GABAergic synapse (*SLC38A3, SLC6A1* and *SLC6A5*), or act as sodium-hydrogen exchangers (*SLC9A8*) or sodium bicarbonate cotransporter in neurons (*SLC4A7* and *SLC4A10*). Finally, *SLC24A4*, a Calcium/Potassium Sodium transporter that is involved in olfactory transduction, hair and eye color, and teeth development. The second cluster of genes is immunity-related, including for example three Toll-like Receptors coding genes (*TLR1, TLR6, TLR10), IRAK4* (interleukin-1 receptor-associated kinase 4) and *CD36* (cluster of differentiation 36).

In Europeans (EUR), we identify 12 biological pathways presenting high levels of introgression, which can be assigned to four non-overlapping clusters of genes (Table 1). Among others, these pathways include “Processing of Capped Intron-Containing Pre-mRNA”, “Uptake and actions of bacterial toxins”, three Netrin-related signaling pathways, and two olfactory signaling pathways. The first cluster of genes obtained by pooling connected subnetworks contains 24 olfactory receptors. The second group of genes is related to hormone signaling, with genes such as *ESRRB* and *ESRRG* (Estrogen-Related Receptors Beta and Gamma), *PGR* (Progesterone Receptor). The third one is involved in responses to bacterial infection and contains, among others, four genes of the SLC family. The fourth cluster also includes genes involved in response to pathogens such as the 2’-5’-oligoadenylate synthetase family (*OAS1-3*), involved in antiviral response (Drappier and Michiels 2015), the signal transducer and activator of transcription 1 (*STAT1*) and the tripartite motif containing 2 (*TRIM2*), both involved in response to different types of pathogens (Najjar and Fagard 2010; Sarute, et al. 2019).

Finally, in Melanesians (PNG), gene networks enriched in archaic introgression are mostly linked to immunity and metabolism. Five pathways belonging to two clusters contain a significant gene subnetwork: “Uptake and actions of bacterial toxins”, “Antigen Presentation: Folding, assembly and peptide loading of class I MHC”, “Vitamin digestion and absorption”, “Ligand-gated ion channel transport”, “Neurotransmitter uptake and metabolism In glial cells”. The cluster “Uptake and actions of bacterial toxins” is obviously involved in immunity and consists mostly in genes of the SLC family. One of them is also found in EAS (*SLC4A10*), but the others are only found in Melanesians: a sodium/calcium exchanger (*SLC24A3*), a sodium/proton exchanger (*SLC9A9*), and two glucose transporters (*SLC2A2* and *SLC2A12*). Interestingly, this cluster contains another gene related to glucose metabolism: Glucose-6-phosphatase 2 (*G6PC2*) as well as *SPTB, which* encodes the Spectrin-beta protein that plays a role in erythrocyte membrane stability. The cluster “Antigen Presentation: Folding, assembly and peptide loading of class I MHC” contains growth factors and immunity-related genes. Cell proliferation related genes include the Growth Hormone Receptor (GHR), Epidermal Growth Factor (*EGF*), and Fibroblast Growth Factor 10 (*FGF10*). We also identify growth factor receptors such as Receptor tyrosine-protein kinase erbB-3 (*ERBB3*), LIF Receptor Alpha (*LIFR*). Metabolism-related genes include kinase suppressor of ras 2 (*KSR2*) that is involved in glucose and fatty acid oxidation. The other genes of this cluster are involved in immune response to pathogens: Janus Kinase 1 (*JAK1*), Mitogen-activated protein kinases (*MAPK1, MAP2K1, MAP2K2*), as well as three Interleukins receptors (*IL20RA, IL22RA2, OSMR*).

We decided to have a closer look at the Spectrin-beta (SPTB) gene, which codes for a protein with a specific function in erythrocyte membrane structure, as it is a constituent of the cytoskeletal network underlying the membrane. This gene is interesting as it could be potentially be involved in response to malaria, a disease that is present in Papua-New-Guinea. As we did not have access to the genotypes of the individuals used for inferring the introgression map, we examined the genomes of 14 Papuan individuals from the Simons Genome Diversity Panel (SGDP) (Mallick, et al. 2016). The comparison of the genotypes of modern individuals to those of three archaic individuals allowed us to confirm the presence of an introgressed haplotype that segregates in Australo-Melanesian populations (Figure 1). The introgressed haplotype overlaps with the SPTB gene and its introgression frequency is maximum in the 5’ promoter region (Figure 1), suggesting that gene expression is affected. Among SGDP samples, we find one homozygous and five heterozygous individuals for the introgressed haplotype, the 15 others carry haplotypes that are distinct from archaic sequences (Figure 1). To confirm the role of SPTB variants in erythrocyte membrane stability, we used ClinVar (Landrum, et al. 2015) records to check whether introgressed archaic variants could have an impact on the red blood cell membrane. We identified 15 mutations that were present in homozygous derived state in archaic individuals and in the introgressed haplotype along this gene. Interestingly, these mutations are all linked to elliptocytosis and spherocytosis, and thus involved in an abnormal shape of red blood cells (Table S1).

**Figure 1:**
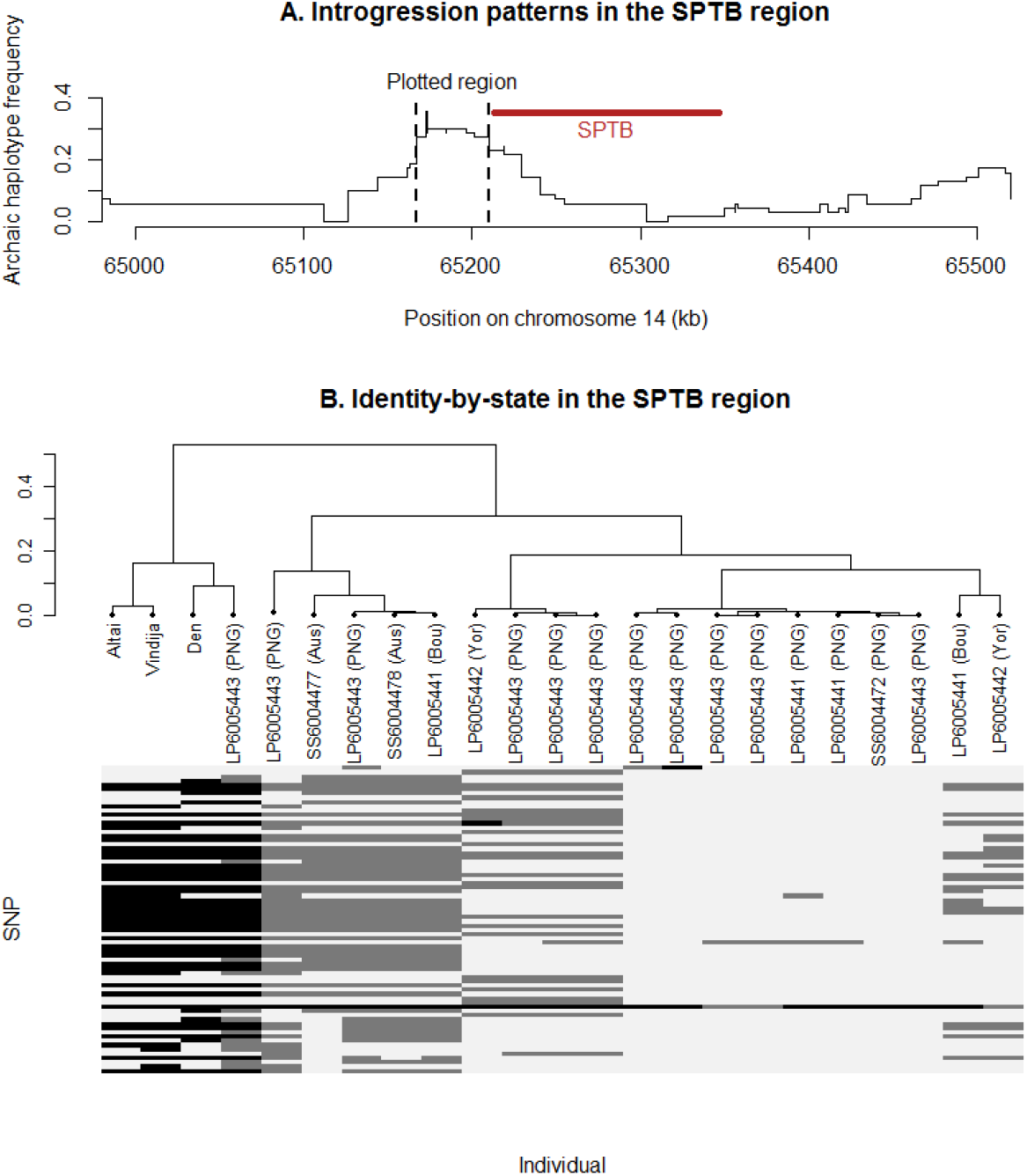
Introgression patterns in the SPTB region. A. Archaic (classified as Neandertal, Denisova or ambiguous) haplotype frequency in the SPTB region based on Vernot et al.’s (2016) introgression map (35 individuals from West New Britain). The location of the SPTB gene is indicated by the red line, and the region used in pane B (with introgression frequency > 0.25) is delimited by vertical dashed lines. B. Relationship between haplotypes found in the SPTB region in SGDP samples from Papua-New-Guinea (PNG), Bougainville (Bou), Australia (Aus), Yoruba (Yor) and archaic individuals (Altai and Vindija Neandertals and Denisova). A neighbor-joining tree based on identity-by-state in the SPTB region is represented on the upper part of pane B. The lower part of pane B corresponds to the genotypes of these individuals for SNPs in this region that have no missing data and a minor allele frequency > 0.05 (grey: heterozygous; black: homozygous for the alternate allele).

### Detection of co-introgressed gene sets

#### Analysis of highly-introgressed gene networks

Previous results suggest that some functionally related sets of genes present an overall enrichment in archaic alleles. We introduce here a new method to test if some introgressed alleles in different genes of a given sub-network present a significant association within individuals, i.e. are in statistical linkage disequilibrium (LD). In practice, we check if the total number of archaic variants within individuals fits a Poisson-binomial distribution (cf. Material and Methods). A significant deviation from expectations indicates the presence of LD (potentially maintained by epistasis) within the gene set. A permutation test then allows us to correct for physical proximity, as we want to focus on long-range LD only.

We applied this new test to all genes belonging to significant subnetworks (Table 1), for each subpopulation separately (i.e. 5 European, 5 East Asian and 1 Papua-New-Guinean subpopulations), such as to avoid detecting spurious signals of LD due to population structure. The test detected significant associations in all 11 populations (one test per population, *p* < 1.10^−5^) without considering physical linkage. However, when using the permutation procedure that considers all genes within 1 Mb as a single locus, the signal disappeared in all gene sets but one (all *p*-values > 0.05). The only remaining significant gene set involves olfactory receptors (ORs) (European cluster 1 in Table 1). These results suggest that i) most of the linkage disequilibrium patterns observed among these genes are due to physical linkage, and ii) that a large (> 1 Mb) archaic haplotype encompassing olfactory receptors is segregating in Europeans.

#### Analysis of the whole pathway database

We then applied the same method to all biological pathways, irrespective of them being found significant by our sub-network analysis or not. With this approach, we find a total of 21 significant pathways. The most significant pathways are the *FMLP Pathway* in East Asians and *SRP Dependent Cotranslational Protein Targeting To Membrane* in Europeans (Figure 2). Papuans show the strongest signal in the *Porphyrin And Chlorophyll Metabolism* (Figure 2). Porphyrins are part of the heme complex and are therefore important hemoprotein components. They are involved in energy metabolism (respiratory chain) and iron and oxygen binding in red blood cells (hemoglobin) and muscles (myoglobin). Other populations present weaker signals of co-introgression. Targeted functions in Europeans are for example related to immunity: *Signaling By TGF Beta Receptor Complex* in CEU, *P75 NTR Receptor Mediated Signaling* in FIN, *Adaptive Immune System, ATF2 Pathway* and *JAK STAT Signaling Pathway* in IBS, and *B Cell Receptor Signaling Pathway* in TSI (Table 2). Some metabolic functions are also significant: *Transport of Vitamins Nucleosides and Related Molecules* in GBR, *Metabolism of Amino Acids and Derivatives* in IBS (Table 2). In East Asian populations, we identify different pathways carrying co-introgressed variants: 6 out of 8 are related to immunity, such as *PI3K events in ERBB4 signaling* in KHV, *AR* and *AT1R pathways* in CHB and CHS, *Factors Involved in Megakaryocyte Development and Platelet Production* in CHS (Table 2). The discrepancies between observed and expected distributions of the sum of archaic alleles per individual are almost always due to an excess of individuals carrying both a too low and a too large number of introgressed alleles (Figure S1). Note that these results barely change when taking a smaller distance threshold of 200 kb. Only two more pathways are significant at a FDR level of 0.05 and they have similar gene contents and functions as the others: *Toll-like receptor signaling pathway* in CEU and *Innate immune response* in CHB.

**Figure 2:**
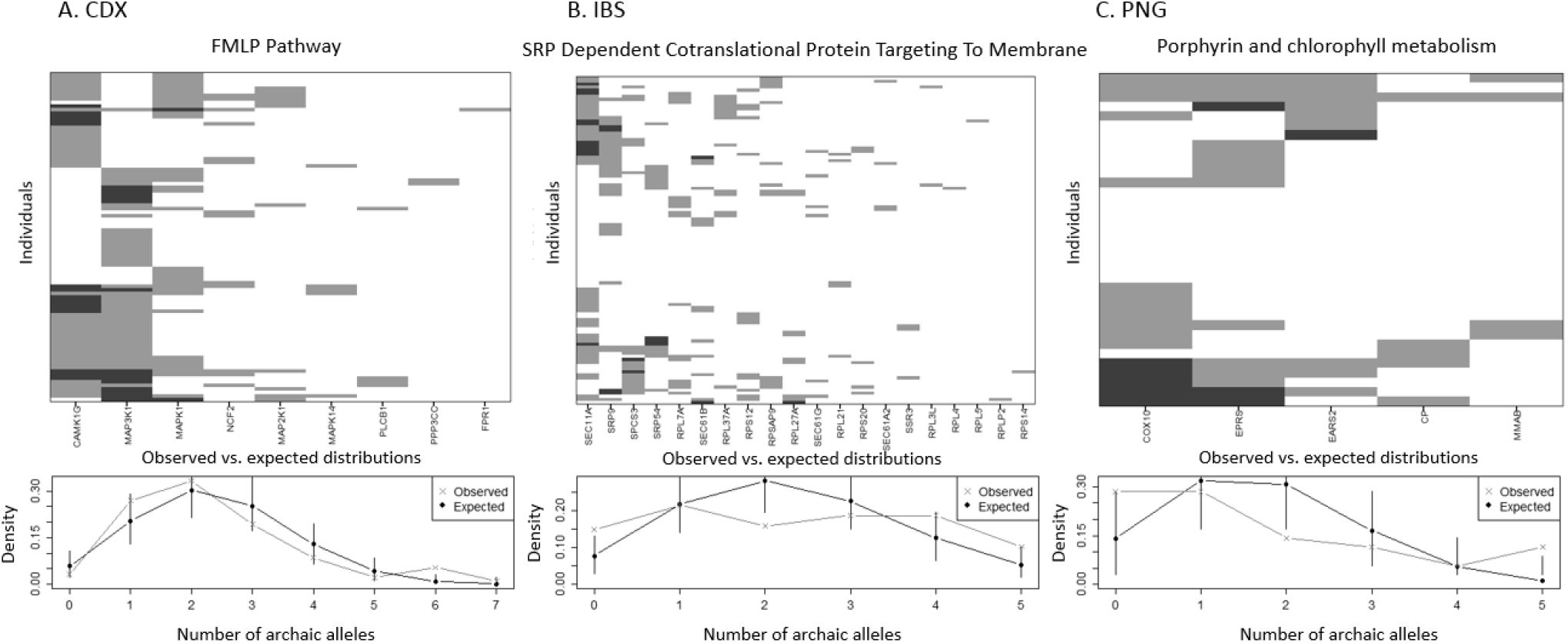
Most significant gene sets with a pattern of co-introgression of archaic variants within individuals in East Asians (A), Europeans (B) and Papua-New-Guineas (C). In each case, the upper part is a heatmap of the number of archaic alleles found within individual for each gene of the pathway (white 0, grey 1, black 2). Each row corresponds to an individual. Individuals (rows) are ordered according to introgression similarity between genes, and genes (columns) are ordered according to introgression frequency. The lower part represents the observed (grey) and expected (black) distributions of the number of archaic variants per individual. Vertical bars delimit 95% confidence intervals of the Poisson-binomial distribution.

**Table 2:**
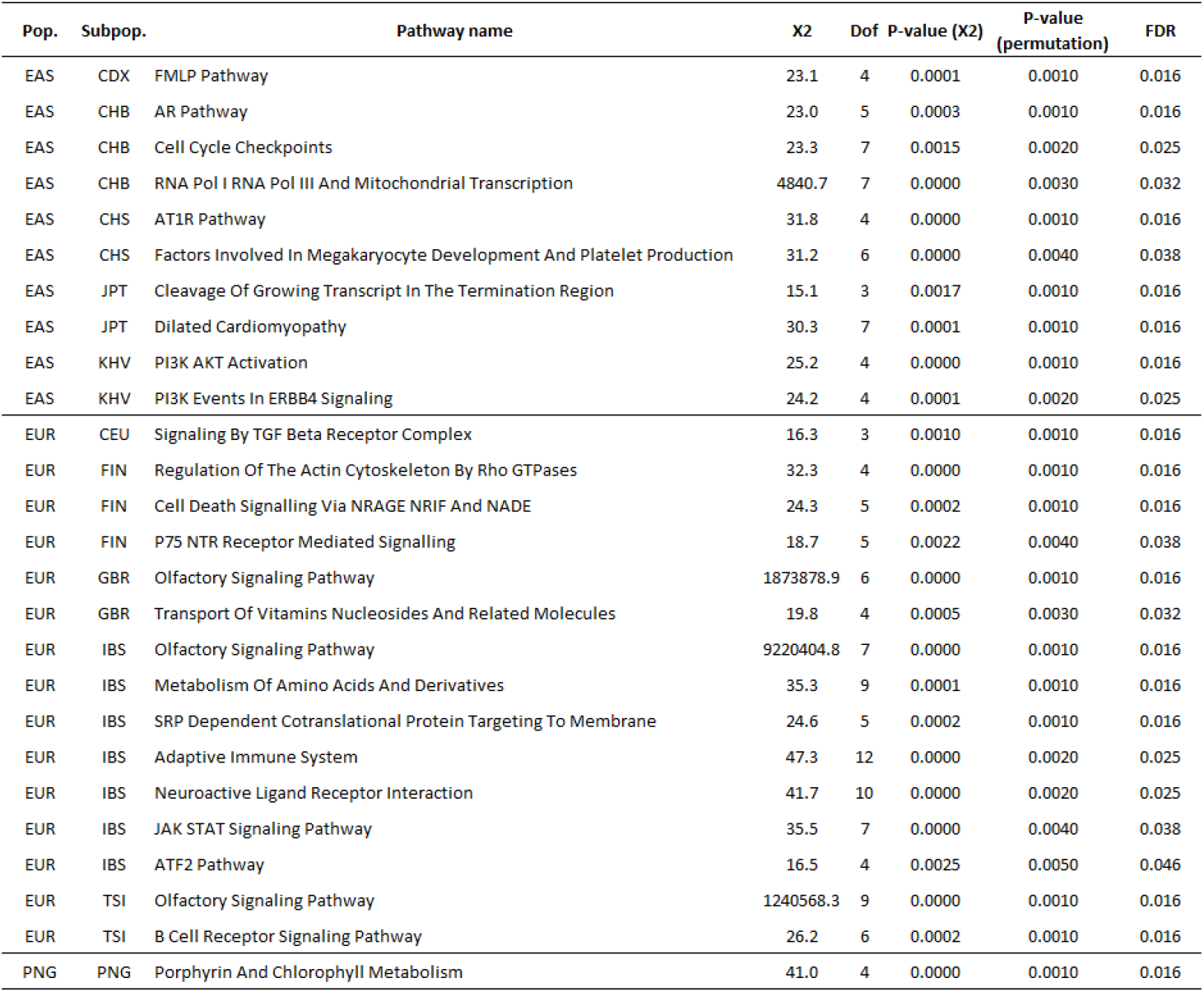
Significant gene sets for the co-introgression test. For each population and subpopulation, significant (FDR < 0.05) pathways names are reported, as well as the χ^2^ value, degrees of freedom and p-values of the goodness-of-fit test. P-values of the permutation test considering linkage between genes are also indicated, as well as the FDR obtained after a Benjamini-Hochberg correction.

| Pop. | Subpop. | Pathway name | X2 | Dof | P-value (X2) | P-value (permutation) | FDR |
| --- | --- | --- | --- | --- | --- | --- | --- |
| EAS | CDX | FMLP Pathway | 23.1 | 4 | 0.0001 | 0.0010 | 0.016 |
| EAS | CHB | AR Pathway | 23.0 | 5 | 0.0003 | 0.0010 | 0.016 |
| EAS | CHB | Cell Cycle Checkpoints | 23.3 | 7 | 0.0015 | 0.0020 | 0.025 |
| EAS | CHB | RNA Pol I RNA Pol III And Mitochondrial Transcription | 4840.7 | 7 | 0.0000 | 0.0030 | 0.032 |
| EAS | CHS | AT1R Pathway | 31.8 | 4 | 0.0000 | 0.0010 | 0.016 |
| EAS | CHS | Factors Involved In Megakaryocyte Development And Platelet Production | 31.2 | 6 | 0.0000 | 0.0040 | 0.038 |
| EAS | JPT | Cleavage Of Growing Transcript In The Termination Region | 15.1 | 3 | 0.0017 | 0.0010 | 0.016 |
| EAS | JPT | Dilated Cardiomyopathy | 30.3 | 7 | 0.0001 | 0.0010 | 0.016 |
| EAS | KHV | PI3K AKT Activation | 25.2 | 4 | 0.0000 | 0.0010 | 0.016 |
| EAS | KHV | PI3K Events In ERBB4 Signaling | 24.2 | 4 | 0.0001 | 0.0020 | 0.025 |
| EUR | CEU | Signaling By TGF Beta Receptor Complex | 16.3 | 3 | 0.0010 | 0.0010 | 0.016 |
| EUR | FIN | Regulation Of The Actin Cytoskeleton By Rho GTPases | 32.3 | 4 | 0.0000 | 0.0010 | 0.016 |
| EUR | FIN | Cell Death Signalling Via NRAGE NRIF And NADE | 24.3 | 5 | 0.0002 | 0.0010 | 0.016 |
| EUR | FIN | P75 NTR Receptor Mediated Signalling | 18.7 | 5 | 0.0022 | 0.0040 | 0.038 |
| EUR | GBR | Olfactory Signaling Pathway | 1873878.9 | 6 | 0.0000 | 0.0010 | 0.016 |
| EUR | GBR | Transport Of Vitamins Nucleosides And Related Molecules | 19.8 | 4 | 0.0005 | 0.0030 | 0.032 |
| EUR | IBS | Olfactory Signaling Pathway | 9220404.8 | 7 | 0.0000 | 0.0010 | 0.016 |
| EUR | IBS | Metabolism Of Amino Acids And Derivatives | 35.3 | 9 | 0.0001 | 0.0010 | 0.016 |
| EUR | IBS | SRP Dependent Cotranslational Protein Targeting To Membrane | 24.6 | 5 | 0.0002 | 0.0010 | 0.016 |
| EUR | IBS | Adaptive Immune System | 47.3 | 12 | 0.0000 | 0.0020 | 0.025 |
| EUR | IBS | Neuroactive Ligand Receptor Interaction | 41.7 | 10 | 0.0000 | 0.0020 | 0.025 |
| EUR | IBS | JAK STAT Signaling Pathway | 35.5 | 7 | 0.0000 | 0.0040 | 0.038 |
| EUR | IBS | ATF2 Pathway | 16.5 | 4 | 0.0025 | 0.0050 | 0.046 |
| EUR | TSI | Olfactory Signaling Pathway | 1240568.3 | 9 | 0.0000 | 0.0010 | 0.016 |
| EUR | TSI | B Cell Receptor Signaling Pathway | 26.2 | 6 | 0.0002 | 0.0010 | 0.016 |
| PNG | PNG | Porphyrin And Chlorophyll Metabolism | 41.0 | 4 | 0.0000 | 0.0010 | 0.016 |

### Specific analysis of olfactory receptors (ORs)

The pattern of introgression observed for olfactory receptors in Europeans is intriguing. Indeed, since 24 genes of this pathway carry an archaic variant, which suggest that the introgressed haplotype could be very large. Many ORs are indeed physically close to each other, most of them being relatively small (typically a single exon of about 1 kb), and a large cluster of OR genes is located near the centromere of chromosome 11 (represented in red around position 53.4 Mb in Figure S2). The pattern of archaic introgression in this region is mainly due to the presence of a large archaic haplotype that segregates in European populations. The frequency of this haplotype is about 10 %, and it is located close to the centromere (downstream at position 55-57 Mb, Figure S2).

We then explored the evolutionary relationships between introgressed and non-introgressed haplotypes at this locus, and particularly those haplotypes present in the 1000 Genomes European individuals (phase III), that we compared to haplotypes found in the Altai (Alt) and Vindija (Vin) Neandertals, and that found in the Denisovan individual (Den). We followed the approach of Danneman et al. (2016) to reconstruct haplotype networks (see Material and Methods). The strategy is to identify clusters of haplotypes (*core* haplotypes) based on their genetic similarity. We found a set of six modern human core haplotypes (labelled I-VI) and three archaic haplotypes (labelled VII-IX). We then computed pairwise nucleotide distances between these 9 core haplotypes, from which we build a neighbor-joining tree (Figure 3).

**Figure 3:**
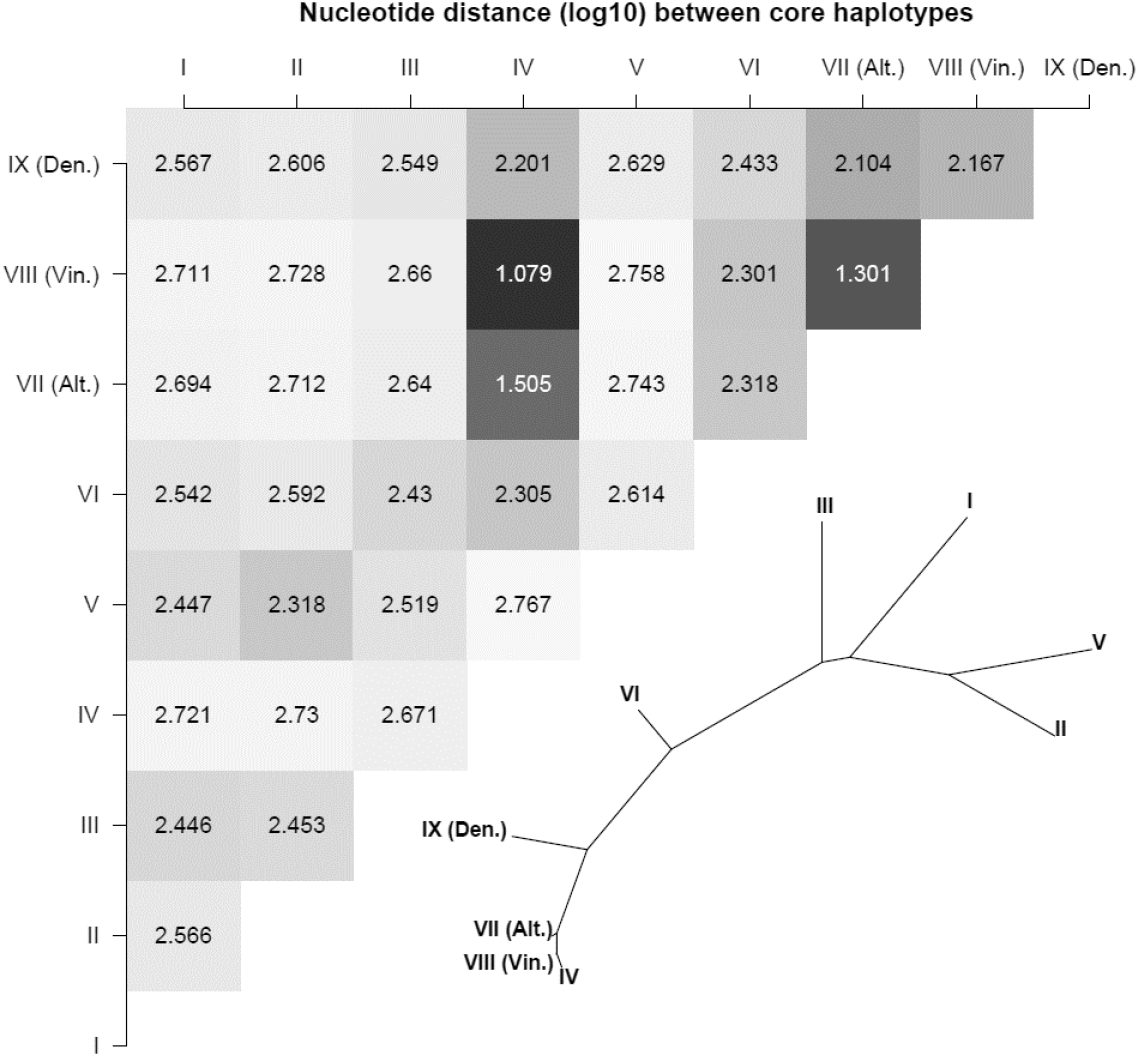
Nucleotide distance (log_10_) between the 9 core haplotypes (I-VI: modern Europeans haplotypes; VII-IX: archaic haplotypes) and neighbor-joining tree based on this matrix.

We see on Figure 3 that haplotype IV carried by modern Europeans is much closer to the archaic haplotypes than all other modern haplotypes. Following Huerta-Sanchez et al. (2014), we computed the probability of observing such a long haplotype by incomplete lineage sorting (ILS) given past demography and recombination rate (see Material and Methods). Even though we are in a very low recombination region, we find that it is highly unlikely to observe such a long haplotype (>= 500 kb) due to ILS (*p* = 1.22.10^−19^) without introgression. This shared ancestry is thus more likely due to an introgression event than to the persistence of an ancestral polymorphism in humans. These observations therefore suggest that this introgression event is recent and happened only in the ancestors of European populations.

## Discussion

### Widespread signal of introgression in immunity-related genes

Several genes involved in immunity carry high-frequency archaic variants. For instance, we identify in East Asians a cluster of five genes, including three Toll-like Receptors (TLR), which show high rates of introgression. These three TLR genes are actually a cluster of physically linked genes, which are known to have been under positive and balancing selection in Eurasians (Dannemann, et al. 2016; Deschamps, et al. 2016; Quach, et al. 2016). We also identify two other TLR-associated genes that have an excess of Neanderthal tracts in East Asians: IRAK4 and MYD88. These two genes are involved in the immune response to pathogens and are functionally linked to TLR receptors as they code for proteins that are activated by TLR to form a protein complex. We also identify three genes with an excess of introgressed variants that all belong to the OAS (2’-5’-oligoadenylate synthetase) cluster in Europeans. These genes are also involved in response to pathogens and have been previously described as candidates for adaptive introgression (Mendez, et al. 2013; Sams, et al. 2016). Another example of introgressed immunity-related genes is the regions containing STAT2 and ERBB3 in Papuans. It has indeed been shown (Mendez, et al. 2012) that a large 250 kb haplotype of Denisovan ancestry segregates at a high frequency and encompasses these genes. Even though we do not identify in our significant networks an excess of introgressed haplotypes in the STAT2/ERBB3 cluster region, we find closely related genes, e.g. JAK1, and numerous other highly introgressed genes that are involved in the immune response (Table 1). It suggests that other functionally related genes might have been targeted by selection, and that adaptive introgression in this case involves more genes than previously thought.

### Possible resistance to malaria

Interestingly, some of the introgression signals we identify in Papuans can be associated to genes potentially involved in response to malaria. A common strategy of resistance to malaria indeed involves defects in normal red blood cells metabolism and structure (Williams 2006). It has been observed that several mutations conferring sickle cell anaemia (haemoglobin S), thalassaemia, Southeast Asian ovalocytosis or Glucose-6-phosphate dehydrogenase (G6PD) deficiency are naturally protective against malaria (reviewed in Williams (2006)). Ovalocytosis, a hereditary condition in which most erythrocytes have an oval shape, is a phenotype that occurs in up to 20% or more in the PNG population (Amato and Booth 1977). Red blood cell membrane defects have several genetic bases, one of them being a mutation in the SPTB gene (Iolascon, et al. 2003; Lelliott, et al. 2017), which is one of our genes candidate for being subject to adaptive introgression. Some mutations in the SPTB gene that are found in archaic genomes and that segregate in Papuans, but which are almost absent from the rest of the world have been previously associated to the elliptocytosis phenotype (see ClinVar records in Table S1). It is also interesting to see that two glucose transporters (GLUT2 and GLUT12) and the G6PD2 gene are also carrying a significant excess of archaic variants. Glucose metabolism in red blood cells has been shown to have a major impact on the survival of *Plasmodium* parasites (Williams 2006), and therefore conditions the susceptibility of humans to malaria. We postulate that introgression could have been the source of several adaptive variants increasing the fitness of individuals in an environment where malaria is endemic. Since malaria resistance alleles typically appeared after the Neolithic transition (Hedrick 2012), these alleles could have been present in archaic populations for a long time, evolving neutrally in the absence of malaria and been positively selected only once the disease appeared (Laval, et al. 2019). Alternatively, archaic populations living in tropical areas could have been exposed to similar endoparasites, which is possible as malaria parasites infect many vertebrates including great apes, which are the most likely origin of the parasite transmission to humans (Prugnolle, et al. 2011). In any case, our results confirm that archaic introgression is widespread in immunity-related genes and that pathogens represent a strong selective pressure which could be one of the major causes of adaptive evolution in humans (Fumagalli, et al. 2011; Daub, et al. 2013; Dannemann, et al. 2016; Quach, et al. 2016).

### Introgression in the solute carrier (SLC) family

Genes from the SLC family encode for membrane transport proteins and are involved in several distinct biological processes. Neandertal variants at two SLC loci (SLC6A11, SLC6A13) have previously been associated to behavioural traits (depression, mood disorders, smoking behaviour) and some alleles have been shown to be preferentially expressed in the brain (Simonti, et al. 2016; Dannemann and Kelso 2017). In Papuans, we find genes showing a significant excess of introgression that have been respectively associated to autism susceptibility and attention-deficit/hyperactivity disorder, e.g. SLC9A9 (Lasky-Su, et al. 2008). Here, we report other genes from the same family that have a brain-biased expression and show an excess of introgressed segments in East Asians and Europeans: SLC6A1 (a GABA transporter), SLC6A5 (a sodium- and chloride-dependent glycine neurotransmitter transporter), and SLC28A1, as well as in Papua-New-Guineans: SLC4A10 (controlling intracellular pH of neurons, the secretion of bicarbonate ions across the choroid plexus, and the pH of the brain extracellular fluid). These results suggest that archaic introgression might have also affected behavioural/neuronal traits, even though it is difficult to link these phenotypes to a precise selective pressure. Other SLC genes showing high levels of introgression include genes encoding for two glucose transporters in Papua-New-Guineans (SLC2A2 and SLC212), and SLC24A4 in East Asians that has been associated to hair and skin pigmentation and adaptive introgression of an archaic variant at this locus has been hypothesized (Dannemann and Kelso 2017). The rest of the SLC genes have less specific functions and are therefore difficult to associate to a particular biological process.

### Olfactory receptors (ORs)

A large Neandertal haplotype encompassing several ORs in the centromeric region of chromosome 11 is found at a frequency of about 10 % in Europeans. A recent study suggests that the whole centromeric region has been introgressed in Europeans (Langley, et al. 2019). These genes are short and clustered together, explaining why ORs pathways are significant in enrichment test that assume independence between loci. Interestingly, more than 40 % of the 856 olfactory receptor genes in the human genome are in 28 gene clusters on chromosome 11 (Taylor, et al. 2006). However, even though the enrichment pattern we observe could be due to physical linkage, we observe a very long haplotype overlapping the centromere of chromosome 11 that includes many olfactory receptors and which has been recently introgressed in humans from Neandertals. Indeed, the recombination rate of this region is very low, but it is not low enough to make it compatible with incomplete lineage sorting since the divergence with archaic humans. However, it is sufficiently low to still observe such a long haplotype tens of thousands years after the introgression event. Even though a case of adaptive introgression is likely for this region, it is difficult to identify the exact selective constraints having promoted this polymorphism in this region, as the relationship between these ORs and a human phenotype remains to be determined.

### A new test to detect co-introgressed variants

In this study, we have introduced a new approach to detect a non-random association of introgressed alleles within individuals at the gene-set level. This approach is similar to a test of linkage disequilibrium but i) it is performed at the individual level rather than at the population level, ii) it is restricted to functionally related, biologically meaningful, sets of genes; and iii) we perform a single test per gene set, irrespective of the number of genes present in this gene-set, which avoids the problem of multiple testing. This approach might thus be a good way to identify small groups of genes that have variants with positive epistatic interactions, which would deserve further investigations and functional validation.

When investigating the genomic signal of polygenic selection from genomic data only, it is difficult to distinguish between two possible scenarios: sequential selection of independent genes or simultaneous selection of alleles at different loci within individuals (through positive epistasis). Provided that alleles can be classified into two categories (introgressed and non-introgressed in our case, our approach allows one to distinguish between these two types of polygenic selection: independent selection at different loci, and epistatic selection within a gene set. By applying this method to archaic introgression in modern humans, we found evidence of these two types of selection. Note however that our test is quite general, as it could be applied to any other distinctions such as two different genetic backgrounds or ancestral/derived states.

Our approach shows that the genomic signal of polygenic selection we observed is most of the time explained by an independent and probably sequential selection of loci involved in related functions (e.g. immune or metabolic functions, Table 1). Our results also highlight the fact that physical linkage might be responsible for a large portion of the signal observed in previous enrichment tests. Indeed, physical linkage between loci will artificially inflate the proportion of significant genes in a given gene set. For example, in the case of archaic introgression, olfactory receptors have been identified several times in enrichment test (Sankararaman, et al. 2014; Steinrücken, et al. 2018), but we observe that highly-introgressed ORs (at least 24 of them) are clustered together in the genome, potentially leading to a spurious signal of enrichment when considering these genes as independent. It follows that one should consider linkage between loci when performing enrichment tests.

Interestingly, we find several significant gene sets with signals of epistatic selection of archaic variants in Europeans, East Asians and Papuans. Most of these gene sets are involved in immunity, and a few are involved in metabolic functions. Even though the observed signal is relatively weak (see Figures 2 and S1), it is still significant after correction for multiple tests. Note that the observation of a strong LD between physically unlinked loci requires extremely strong epistatic selection (Felsenstein 1965), so that it is still remarkable that we observe signals of epistasis given the relatively small samples sizes on which our test was applied.

## Conclusions

Our detailed examination of the distribution of introgressed haplotypes in the human genome has allowed us to identify several genes interacting in biological pathways that carry archaic variants at a relatively high frequency. Overall, our results suggest that archaic introgression has affected human metabolism and response to different types of pathogens (bacteria, virus and protists), which have been critically determinant during human adaptive history. We also propose a new test to distinguish between the independent selection and co-selection of adaptive variants, a distinction that is usually overlooked in classical tests of polygenic selection. Even though the overall amount of Neandertal and Denisovan introgression is quite low in modern humans, some introgressed genes nevertheless present archaic variants that are in linkage disequilibrium within individuals, which is highly unexpected for variants found in different location of the genome without a strong selective force maintaining and possibly promoting these associations. It suggests that these variants have a strong impact on the present fitness of their carriers and that epistatic selection maintains them at relatively high frequency.

## Material and methods

### Genetic data and introgression maps

To study introgressed haplotypes in present-day human genomes, we used introgression maps published in Vernot et al. (Vernot, et al. 2016). This dataset consists in haplotype calls for Neandertal, Denisova in Europe (EUR, 503 diploid individuals), East Asia (EAS, 504 diploid individuals), and Papua-New-Guinea (PNG, 35 diploid individuals). Putatively introgressed regions were detected based on the *S*\* statistic (Wall, et al. 2013). These regions were then compared to the reference Neandertal genome to characterize introgressed archaic haplotypes in modern humans (Vernot, et al. 2016). These introgression maps are available online [https://akeylab.princeton.edu/downloads.html].

To study the functional impacts of archaic variants, we estimated the amount of archaic introgression in each population for each gene. We define the level of introgression for a given region of the genome as the average frequency of inferred archaic variants of this region. To be more conservative, we computed this measure only in the exons of each gene. We considered Neandertal haplotypes only for Europeans and East Asians, and Neandertal and Denisova haplotypes for Papua-New-Guineans. In the end, we have a single introgression measure per gene and per population, to which we will refer below as the *gene score* or *introgression score*.

### Detection of gene subnetworks enriched in archaic variants

#### Biological pathways data and conversion to gene networks

We considered biological pathways as gene networks. More formally, we define a gene network as a graph *G*(*V, E*), where *V* is a set of nodes (i.e. genes), and *E* is a set of edges (i.e. interactions between genes). In this study we used three signaling and metabolic pathway databases that are considered as references in systems biology: (i) KEGG, the Kyoto Encyclopaedia of Genes and Genomes Pathway database; (ii) NCI, the National Cancer Institute / Nature Pathway Interaction Database (Schaefer, et al. 2009); and (iii) Reactome (Fabregat, et al. 2016). We then used the R/Bioconductor graphite package to convert biological pathways into graphs of interacting genes (Sales, et al. 2012).

#### Search for outlier subnetworks

We applied the workflow described in Gouy et al. (2017) and implemented in the R/Bioconductor package *signet* to detect highly introgressed gene subnetworks within biological pathways. This approach uses a simulated annealing algorithm to find the highest-scoring subnetwork (HSS) within each biological pathway. Then, a statistical testing procedure is implemented to detect outlier HSS. An empirical null distribution of HSS scores is built, from which a p-value is then computed.

We applied this procedure to the three populations (EAS, EUR, PNG) for which we computed the exonic introgression score. For each population, we ran the algorithm on each individual pathway. We used 5,000 iterations of simulated annealing for individual pathways. 10,000 permutations of gene scores were done to generate the empirical null distribution and compute p-values.

### Core haplotype analysis

We explored the relationships between modern and archaic haplotypes at the region encompassing olfactory receptors (ORs) significant subnetwork. We took European individuals from the 1000 Genomes phase III dataset (ftp://ftp.1000genomes.ebi.ac.uk/vol1/ftp/release/20130502/), as well as three archaic individuals: the two Altai and Vindija Neandertals, and Denisova. We followed the approach of Danneman et al. (2016) to reconstruct haplotype networks as follows. We used all 980 SNPs that were polymorphic in Europeans in this 5 Mb region and where archaic hominins genomes were homozygous. We merged the resulting haplotypes into core haplotypes that differed by less than 350 nucleotides (i.e., ~ 1/1,000 base pairs mismatch between two chromosomes). This resulted in a set of six modern human core haplotypes (labelled I-VI) and three archaic haplotypes (labelled VII-IX, which are simply the Neandertal and Denisova genome sequences). We generated a consensus sequence for each modern human core haplotype by taking the major allele at each position. We then represented the nucleotide distance matrix between all the nine core haplotypes and we built a neighbor-joining tree based on this distance matrix using the R package *ape* (Paradis, et al. 2004).

### Incomplete lineage sorting analysis

We used the method of Huerta-Sanchez et al. (2014) to compute the probability of observing an archaic haplotype of a given size by incomplete lineage sorting (ILS) given some demographic parameters and recombination rate. Just as in the original study, we obtained the average recombination rate from the three maps available on UCSC (sex-averaged: deCode 0.06 cM/Mb) for this 500kb region. We modelled the expected length of segments of ILS given the local recombination rate, time since the divergence from the common ancestor, and generation time, and compare this to the length of the observed introgressed haplotype. We used a generation time (g) of 25 years and the average recombination rate estimate from UCSC (*r* = 5 × 10^−9^). Estimates of branch lengths since divergence times (in years) are taken from Prüfer et al. (Prüfer, et al. 2017). To be conservative, we used the shortest possible branch lengths (in years) from the estimated ranges because it allows for fewer recombination events, following Dannemann et al. (2016). We computed the expected length *L* for two recent estimates of the human mutation rate; μ = 1 × 10^−9^ and μ = 0.5 × 10^−9^ per base pair per year: 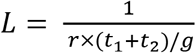, with *g* being the generation time (25 years). The probability of a length of at least *X* kb as 1 – Γ(*X*, shape = 2, rate = 1/*L*) with Γ being the gamma distribution function.

### ClinVar records for Spectrin-B

All ClinVar records associated with the SPTB gene were downloaded from NCBI website (https://www.ncbi.nlm.nih.gov/clinvar/?term=SPTB[gene], on the 04/09/2018). Out of the 178 extracted records, we retained 15 mutations that were found in at least one of the three archaic genomes (Altai Denivosa, Altai and Vindija Neandertals genotypes (Prüfer, et al. 2017) downloaded from http://cdna.eva.mpg.de/neandertal/Vindija/VCF/) using bcftools version 1.5 (Li 2011). We extracted the genotypes of individuals from the Simons Genome Diversity Project (https://www.simonsfoundation.org/simons-genome-diversity-project/) along this region.

### Test of co-introgression in a gene set

Here, we present a new method to detect if some groups of variants (here, archaic haplotypes) are co-occurring more than expected by chance alone in some individuals, or if there is no significant association between loci within individuals. We estimate this measure of *functional linkage* for all considered gene sets (e.g. biological pathways). At each locus, each individual, will have a score of 0, 1 or 2, depending on its genotype, which represents the number of archaic haplotypes observed at this locus. The statistical test consists in comparing, for each gene set, the observed distribution of the sum of archaic haplotypes per individual to its expectation.

#### Co-introgression test

Let *G* be a set of genes. For each gene *g* ∈ *G*, let *x_ig_* ∈ {0,1,2} be the observed number of archaic haplotypes of individual *i* present at gene *g*. Let *p_g_* be the population frequency of archaic haplotypes at a given gene *g*. For a given individual, the expected number of archaic haplotypes is the sum of *K* Bernoulli trials (one trial per homologous chromosome and per gene, i.e. *K* = 2 |*G*| trials). Therefore, we model these observations *x* as a Poisson binomial distribution, which corresponds to a sum of independent Bernoulli trials *that are not identically distributed*, since the probability to have an archaic haplotype varies for each gene. That way, we can model the outcome of different loci for a given individual with locus-specific probabilities of success (i.e. estimated frequencies at this locus).

The probability mass function of the Poisson binomial distribution is:

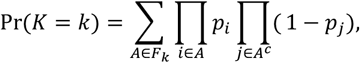

where *F_k_* is the set of all subsets of *k* integers that can be selected from {1,2,3,…, *n*}. *A^c^* is the complement of set *A*. Pr(*K* = *k*) is the sum of the products of success probabilities (allele frequencies) over all possible subsets of size *k*. To compute this sum, one needs to enumerate all elements in *F*_k_, which is not feasible in practice. We used the discrete Fourier transform implemented in the R package poisbinom to estimate it (Hong 2013).

Now that we know the distribution of the sum of archaic alleles per individual, we want to test the goodness-of-fit to a theoretical Poisson binomial (with parameters *n* and *p*_g_). We perform this comparison using a chi-square test, each category being a possible value of the sum of archaic alleles (*k*) per individual.

#### Controlling for physical linkage and population structure

The test we have developed assumes that loci are independent, which might not be true due to population structure or physical linkage. To exclude a gene set presenting linkage because of one of these factors, we apply a permutation test if the exact test is significant. First, we apply this test to subpopulations of the 1000 Genomes project (i.e. for Europeans: CEU, FIN, TSI, IBD, GBR; and East Asians: CDX, CHB, CHB, KHV, JPT). We do that to avoid a spurious signal due to a Wahlund effect due to population structure. Then, we permute the genotypes 10,000 times within gene clusters. We cluster genes based on their pairwise physical distance, using a neighbor-joining tree that we cut at a given distance threshold (here, 1 Mb) to define the clusters. At each iteration, we sample one cluster of gene, permute the genotypes among individuals, and compute the chi-square statistic used to test the goodness-of-fit of the distribution. Finally, the resulting empirical distribution of χ^2^ statistics is used to compute a p-value.

#### Application to real data

We applied this method to the introgression maps from Vernot et al. (Vernot, et al. 2016) mentioned above. One first needs to get, for each gene and each chromosome, the number of introgressed alleles. We generated a matrix of *n* individuals x *g* genes, with entries equal to 0, 1 or 2, depending on the number of archaic allele copies found at the site with the highest archaic allele frequency for a given locus. We then applied the method to different gene sets: i) the significant subnetworks resulting from the simulated annealing analysis; and ii) gene sets from the Molecular Signatures Database (MSigDB v. 6.2), a standard collection of annotated gene sets (retrieved from http://software.broadinstitute.org/gsea/msigdb/ on the 02/03/2019). We analysed the pathway set C2 (curated databases: BIOCARTA, KEGG, NABA, NCI, PID, and Reactome). All the analyses have been carried out using R 3.4.0.

## Acknowledgments

This work was partially supported by a Swiss National Science Foundation (grant number 310030B-166605) to LE. We thank Alexandre Thiéry for his contribution to bioinformatics analyses.

