## Supplementary material for "Polygenic patterns of adaptive introgression in modern humans are mainly shaped by response to pathogens"

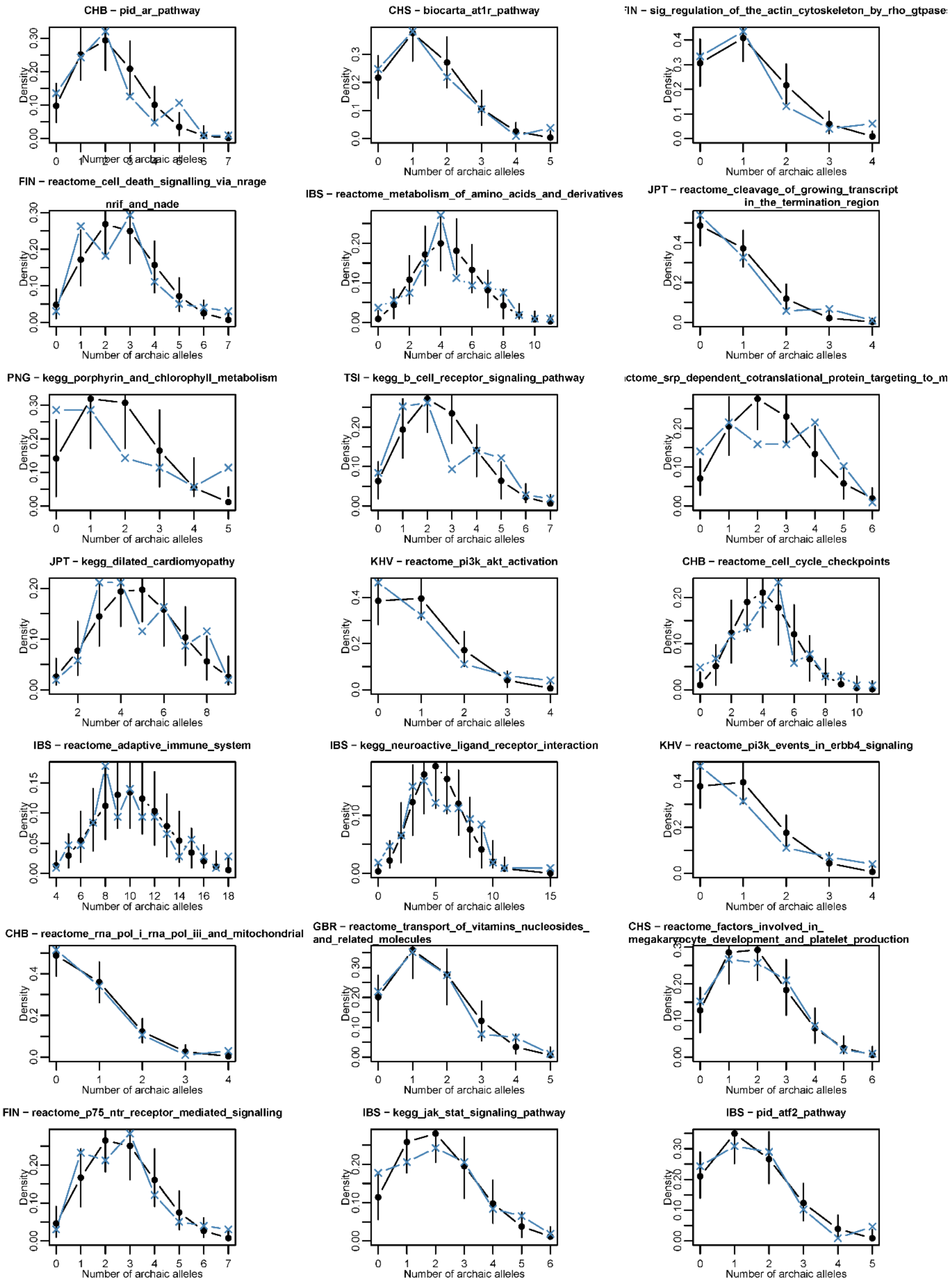


**Figure S1**: Distribution of the number of archaic alleles per individual in pathways significant for the co-introgression test.

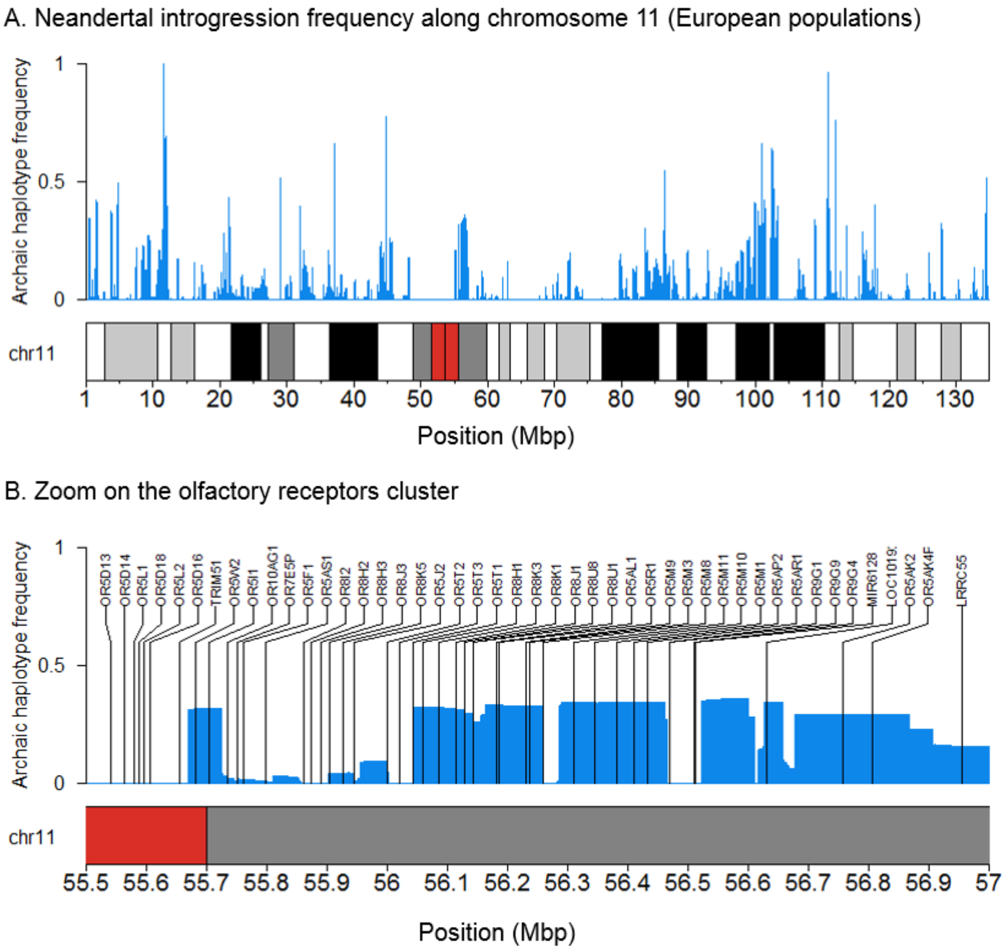


**Figure S2**: Distribution of the frequency of Neandertal variants in Europeans along chromosome 11 (A) and in the olfactory receptors region (B). The upper part of each pane corresponds to the archaic haplotype frequency, the lower part corresponds to the chromosome 11 cytobands (red = centromeric region). The zoomed region is located between positions 55.5 to 57 Mb. The genes found in this region (mostly olfactory receptors) are also indicated.

**Table S1**: ClinVar records for 15 variants in the SPTB gene.

| Name | Condition(s) | Clinical significance | Chr. (GRCh37) | Location (GRCh37) | Den. | Vin. Ne. | Alt. Ne. |
|---|---|---|---|---|---|---|---|
| NM_001355436.1(SPTB):c.6345+321A>G | Elliptocytosis\|Spherocytosis, Dominant | Likely benign(Last reviewed: Jun 14, 2016) | 14 | 65233123 | TRUE | TRUE | TRUE |
| NM_001355436.1(SPTB):c.5943C>T (p.Arg1981=) | not specified\|Elliptocytosis\|Spherocytosis, Dominant | Uncertain significance(Last reviewed: Jun 14, 2016) | 14 | 65235831 | FALSE | TRUE | TRUE |
| NM_001355437.1(SPTB):c.5535C>T (p.Leu1845=) | not specified\|Elliptocytosis\|Spherocytosis, Dominant | Benign/Likely benign(Last reviewed: Apr 13, 2017) | 14 | 65239316 | TRUE | TRUE | TRUE |
| NM_001355436.1(SPTB):c.4860T>C (p.Ile1620=) | not specified\|Elliptocytosis\|Spherocytosis, Dominant | Benign/Likely benign(Last reviewed: Apr 11, 2017) | 14 | 65241228 | TRUE | TRUE | TRUE |
| NM_001355436.1(SPTB):c.4818C>T (p.Tyr1606=) | not specified\|Elliptocytosis\|Spherocytosis, Dominant | Benign/Likely benign(Last reviewed: Apr 13, 2017) | 14 | 65241867 | TRUE | TRUE | TRUE |
| NM_001355437.1(SPTB):c.4779A>G (p.Ala1593=) | not specified\|Elliptocytosis\|Spherocytosis, Dominant | Benign/Likely benign(Last reviewed: Apr 13, 2017) | 14 | 65241906 | TRUE | TRUE | TRUE |
| NM_001355437.1(SPTB):c.4752C>T (p.Asn1584=) | not specified\|Elliptocytosis\|Spherocytosis, Dominant | Benign/Likely benign(Last reviewed: Apr 13, 2017) | 14 | 65241933 | TRUE | TRUE | TRUE |
| NM_001355436.1(SPTB):c.4564-4G>A | not specified\|Elliptocytosis\|Spherocytosis, Dominant | Benign/Likely benign(Last reviewed: May 15, 2018) | 14 | 65242125 | TRUE | TRUE | TRUE |
| NM_001355437.1(SPTB):c.4293A>G (p.Arg1431=) | not specified\|Elliptocytosis\|Spherocytosis, Dominant | Benign/Likely benign(Last reviewed: Apr 13, 2017) | 14 | 65246623 | TRUE | TRUE | TRUE |
| NM_001355436.1(SPTB):c.4208G>A (p.Arg1403Gln) | not specified\|Elliptocytosis\|Spherocytosis, Dominant | Conflicting interpretations(Last reviewed: Feb 28, 2017) | 14 | 65249066 | TRUE | TRUE | TRUE |
| NM_001355436.1(SPTB):c.4003-12T>C | not specified\|Elliptocytosis\|Spherocytosis, Dominant | Likely benign(Last reviewed: Jun 14, 2016) | 14 | 65249283 | TRUE | TRUE | TRUE |
| NM_001355437.1(SPTB):c.3451A>G (p.Asn1151Asp) | not specified\|Elliptocytosis\|Spherocytosis, Dominant | Benign/Likely benign(Last reviewed: Apr 13, 2017) | 14 | 65253232 | TRUE | FALSE | FALSE |
| NM_001355436.1(SPTB):c.2154A>C (p.Ile718=) | not specified\|Elliptocytosis\|Spherocytosis, Dominant | Benign/Likely benign(Last reviewed: Apr 13, 2017) | 14 | 65260227 | TRUE | TRUE | TRUE |
| NM_001355437.1(SPTB):c.1316G>A (p.Ser439Asn) | not specified\|Elliptocytosis\|Spherocytosis, Dominant | Benign/Likely benign(Last reviewed: Apr 13, 2017) | 14 | 65263300 | TRUE | TRUE | TRUE |
| NM_001355436.1(SPTB):c.300+7T>C | not specified\|Elliptocytosis\|Spherocytosis, Dominant | Benign/Likely benign(Last reviewed: Apr 13, 2017) | 14 | 65271650 | TRUE | TRUE | TRUE |
